# A Fast and Memory-Efficient Implementation of the Transfer Bootstrap

**DOI:** 10.1101/734848

**Authors:** Sarah Lutteropp, Alexey M. Kozlov, Alexandros Stamatakis

## Abstract

Recently, Lemoine *et al*. suggested the Transfer Bootstrap Expectation (TBE) branch support metric as an alternative to classical phylogenetic bootstrap support metric on taxon-rich datasets. However, the original TBE implementation in the `booster` tool is compute- and memory-intensive. Therefore, we developed a fast and memory-efficient TBE implementation. We improved upon the original algorithm described by Lemoine *et al*. by introducing multiple algorithmic and technical optimizations. On empirical as well as on random tree sets with varying taxon counts, our implementation is up to 480 times faster than `booster`. Furthermore, it only requires memory that is linear in the number of taxa, which leads to 10× - 40× memory savings compared to `booster`. Our implementation has been partially integrated into `pll-modules` and `RAxML-NG` and is available under the GNU Affero General Public License v3.0 at https://github.com/ddarriba/pll-modules and https://github.com/amkozlov/raxml-ng. The parallelized version that also computes additional TBE-related statistics is available in `pll-modules` and `RAxML-NG` forks at: https://github.com/lutteropp/pll-modules/tree/tbe and https://github.com/lutteropp/raxml-ng/tree/tbe.

## 1 Introduction

The Felsenstein bootstrap (FBP) [2] procedure is widely used to assess the robustness of phylogenies. The FBP draws columns from the multiple sequence alignment (MSA) with replacement 100 or more times (for a discussion of the appropriate number of replicates see Pattengale *et al*. [9]) to generate MSA replicates. Then, for each MSA replicate a corresponding bootstrap (BS) replicate tree is inferred.

Each branch in a phylogenetic tree induces a bipartition (also called *split*) of the set of tips into two subsets. The smaller of these two sets is referred to as the ‘light side’ of the bipartition, whereas the larger set is called the ‘heavy side’. In case both sets are of equal size, the ‘light side’ is chosen arbitrarily.

Subsequently, the bootstrap support value of a branch in the reference tree (e.g., the best-known ML tree on the original MSA) is computed by counting how many BS replicate trees contain the same branch (or respective *bipartition*/*split*), and dividing this count by the total number of BS trees. In the classical FBP approach, only bipartitions that match *exactly* are counted. Conversely, the Transfer Bootstrap Expectation (TBE) metric [6] also takes into account all ‘similar’ bipartitions in the BS replicate trees. The contribution of such similar bipartitions is weighted by their similarity to the respective reference bipartition.

The computation of TBE support is based on the so-called transfer distance. The transfer distance *δ*(*b, b*\*) between a branch *b* in the reference tree and a branch *b*\* in a BS replicate is the minimum number of taxa that need to be moved to transform the bipartition induced by *b* into the bipartition induced by *b*\*.

The transfer index *ϕ*(*b, T*\*) is defined as the minimum transfer distance between a branch *b* in the reference tree and the branches in the BS replicate tree *T*\*:

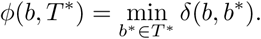

Given a reference tree and a set of BS replicate trees, Lemoine *et al*. define the TBE(*b*) of a branch *b* in the reference tree as follows:

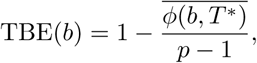

where 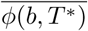 is the average transfer index over all BS replicates and *p* is the number of taxa on the ‘light’ side of the bipartition induced by *b*.

## 2 Implementation

We implemented the transfer bootstrap computation as part of the `pll-modules` library. The `pll-modules` library offers high-level modules for the low-level phylogenetic likelihood library `libpll` [3], e.g., to perform model parameter optimization, tree moves, MSA validation etc.. Besides likelihood computations, `libpll` and `pll-modules` libraries provide highly efficient tree operations such as NEWICK parsing/writing, conducting tree traversals, and manipulating bipartitions. Hence, using `pll-modules` allowed us to leverage these routines for our TBE implementation, and to facilitate integration into third-party programs as well as into our `RAxML-NG` [5] software. Initially, in `RAxML-NG v0.7`, we had implemented a naïve TBE computation method which relied on calculating distances between the reference bipartition and *all* BS tree bipartitions. Despite having a higher theoretical run time complexity of *O*(*n*^2^\**m*) in comparison to *O*(*n*\**m*) [6] (where *n* is the number of taxa and *m* is the number of BS trees), this implementation was faster than `booster` in practice (see Figure 1). However, it scaled poorly to large numbers of taxa. Therefore, we designed and integrated a more efficient implementation into `RAxML-NG v0.8.1` and later versions which we describe in the following.

**Fig. 1.**
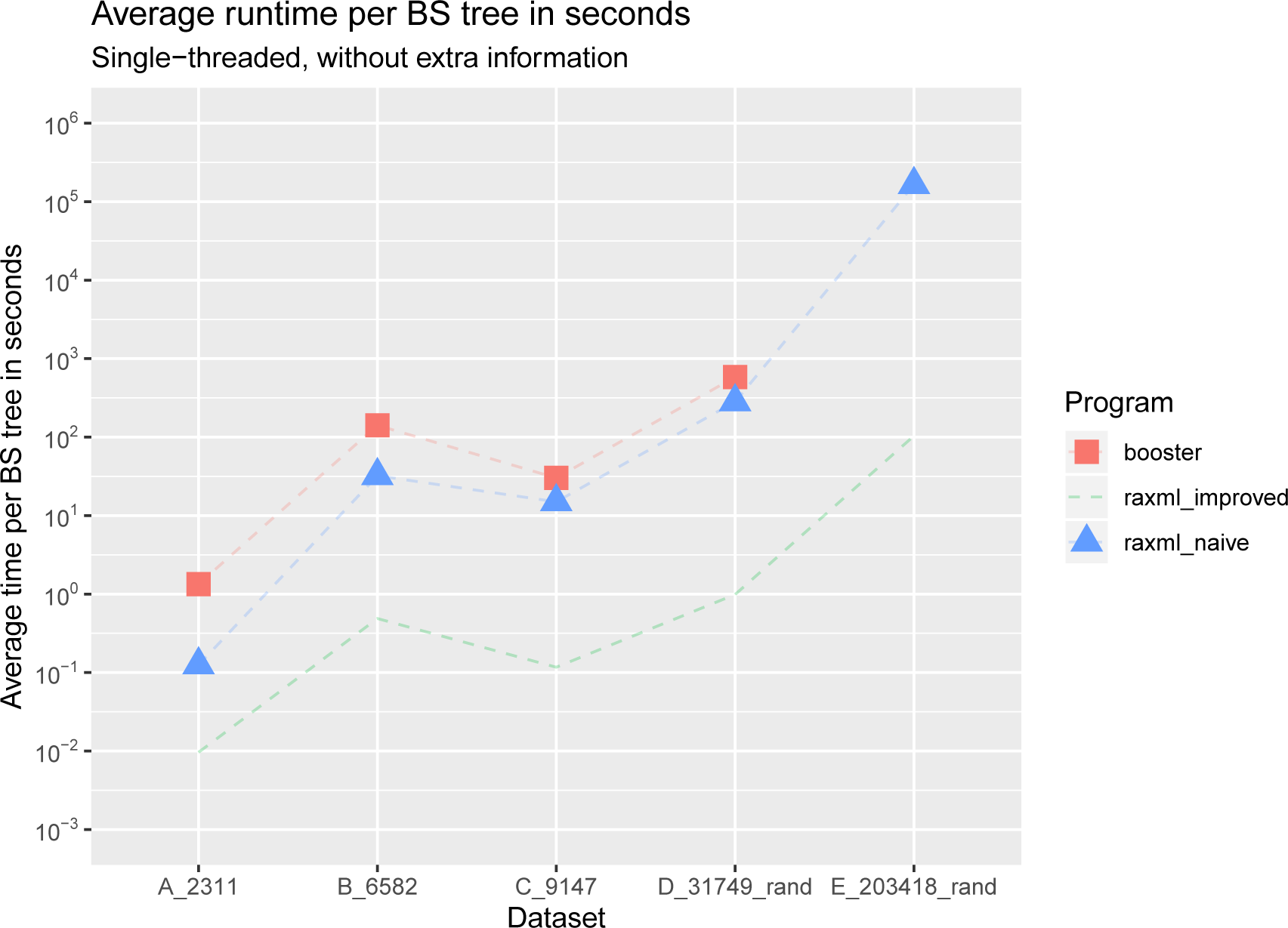
Average runtime per BS tree in seconds, without computing additional information. All tools were executed sequentially. Note the logarithmic scale on the y-axis. On the E_203418_rand dataset, `booster` went out of memory. We can see on this plot that `RAxML-NG improved` is several orders of magnitude faster than `booster` and `RAxML-NG naïve`. The slowest tool across all tested datasets is `booster`.

As in [6], we encode each bipartition as a bit vector.

We compute the transfer index *ϕ*(*b, T*\*) via a post-order traversal as described in [6]. However, we first test if two bipartitions are identical prior to computing the transfer distance, by hashing the bipartitions. We also stop the traversal of *T*\* early if the lowest possible transfer distance (which is *p* − 1) has already been encountered.

The `booster` tool parallelizes the computation *across* BS trees. Our implementation uses OpenMP to parallelize computations *within* each BS tree, as we parallelize over branches in the reference tree for each BS tree. This approach is more fine-grained than the approach followed by booster.

Our per-branch-parallelization has the following advantages (+) and disadvantages (-) compared to the per-tree-parallelization approach used in `booster`:

– (+) less additional memory is required (as only one tree is processed at a time)
– (+) better expected performance in terms of parallel efficiency for high numbers of taxa and low numbers of BS trees
– (-) worse expected performance for low numbers of taxa and high numbers of BS trees

However, as we show in Section 3, our sequential implementation can already process extremely large datasets in but a few minutes.

Our implementation can also print additional per-taxon and per-branch statistics (`-r` and `-c` options in `booster`), and these are also computed more efficiently than in `booster` (see Section 2.4). At present, the OpenMP parallelization and the computation of those additional statistics are not part of the production release of `RAxML-NG`. Our extended version is available in a separate repository at https://github.com/lutteropp/raxml-ng/tree/tbe.

### 2.1 Naïve *O*(*n*^*2*^ * *m*) algorithm (used in RAxML-NG v0.7)

The transfer distance *δ*(*b, b*\*) between two bipartitions *b* and *b*\* can be naïvely computed in linear time via two Hamming distance computations and one minimum operation: For each bipartition, we assign the value 0 to the taxa on one side of the bipartition and the value 1 to the remaining taxa. Then, the transfer distance between *b* and *b*\* is the smaller of the two hamming distances between the bit-vectors induced by *b* and *b*\* and between 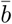 and *b*\*, where in 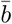 the zeros and ones are swapped.

Using this approach, computing the transfer index *ϕ*(*b, T*\*) requires *O*(*n*^2^) time. Algorithm 1 shows the computation of transfer BS support using our naïve approach.

#### Algorithm 1: Naïve *O*(*n*^2^ * *m*) Algorithm for computing transfer bootstrap support.

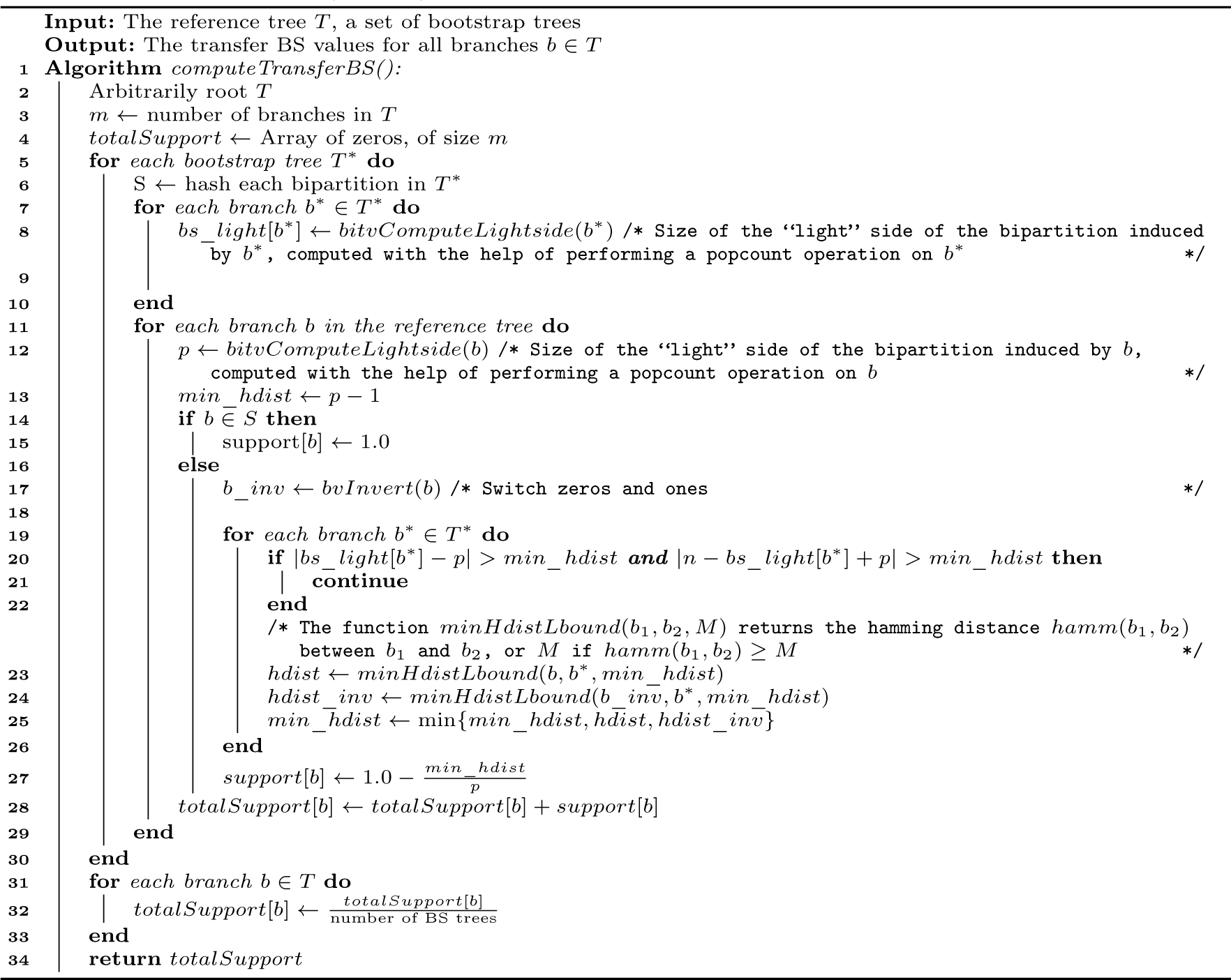

### 2.2 Improved *O*(*n* * *m*) algorithm (used in RAxML-NG v0.8.1 and later)

Let *n* be the number of taxa in the reference tree. We assume that the *m* BS trees share the same set of taxa as the reference tree. Lemoine *et al*. describe the following *O*(*n*) algorithm for computing the transfer index *ϕ*(*b, T*\*) for a branch *b* ∈ *T* and a BS tree *T*\*:

– Let *p* be the size of the “light side” of the bipartition induced by *b*. We assign the value 0 to each taxon on the light side of *b*, and the value 1 to each taxon on the heavy side of *b*.
– Apply the same taxon-value-assignments to the taxa in *T*\*.
– Perform a post-order traversal of *T*\*, counting the number of ones #*ones*_*subtree* in every subtree rooted at a branch *b*\* ∈*T*\*.
– Besides #*ones*_*subtree*, we also know the following values:
  - #*zeros*_*subtree* = *n -* #*ones*_*subtree*
  - #*zeros*_*total* = *p*
  - #*ones*_*total* = *n*− *p* For making the bipartitions *b* and *b*\* identical, we need to either
  - move all zeros inside the subtree and all ones outside, or
  - move all ones inside the subtree and all zeros outside.

Hence, the transfer distance *δ*(*b, b*\*) can be computed as follows:

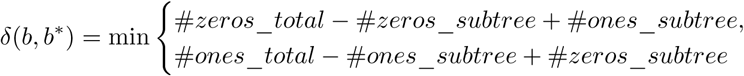

We compute the transfer distance *δ*(*b, b*\*) in *O*(1) during the post-order traversal, using the formula given above. The transfer index *ϕ*(*b, T*\*) is the minimum of the computed transfer distances.

For computing TBE support over *m* BS trees, the overall worst-case complexity of this algorithm is *O*(*n* * *m*).

*Comparison to the algorithm implemented in booster* After inspecting the source code, we realized that `booster` does not implement the algorithm described by Lemoine *et al*. [6], *but instead the algorithm described by Brehelin et al*. [1]. *However, the algorithm by Brehelin et al*. requires quadratic instead of linear memory.

### 2.3 Preprocessing and Speed Improvements

In order to reduce the total number of operations, we preprocess both, the reference tree, and the BS trees. We arbitrarily root the trees at an inner node to have a direction for the post-order traversals needed for computing the transfer index.

Algorithm 2 shows the computation of TBE support using the improved approach, together with our preprocessing and early-stop improvements explained in the following.

#### Preprocessing the Reference Tree

For each branch *b* in the reference tree, we precompute the size *p* of its “light side” via a post-order traversal. Following the description from Lemoine *et al*., during the transfer distance computation we will assign the value 0 to all taxa on the “light side” of the split induced by *b* and 1 to the remaining taxa. To conduct this efficiently, in the preprocessing step, we note for each branch whether the value 0 will be assigned to the taxa within the subtree or the taxa outside the subtree. We also store the index of the leftmost and the index of the rightmost taxon in the subtree. Thereby, we ensure beforehand that the taxa are indexed by 0 to *n*−1 from the left of the tree to the right of the tree. This enables us to efficiently initialize the array which keeps track of the number of ones in the subtrees.

Hence, for each node *v* in the reference tree *T*, we store:

–The indices of the leftmost and rightmost node in the subtree *T*_*v*_ rooted at *v*.
–The size of the “light side” *p*, which is min {|*T*_*v*_|, *n* −|*T*_*v*_ |}.
–A boolean flag whether the taxa in *T*_*v*_ are all 0 or all 1 in the bipartition induced by the incoming edge of *v*.

#### Preprocessing the Bootstrap Trees

As already mentioned, we process the BS trees one after another. Since the subtree sizes in a BS tree do not change for different reference bipartition queries, we precompute the subtree sizes once via a post-order traversal of the BS tree. For each node *v*\* in a BS tree *T*\*, we thus store the size 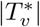 of the subtree 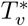 rooted at *v*\*. Moreover, we hash the bipartitions induced by the BS tree branches, such that we can skip the *O*(*n*) transfer index computation for each reference tree bipartition that is also present in the BS tree.

#### Early-Stop

We can easily detect whether a split *b* from the reference tree is present in a BS tree *T*\* via our hashing. In this case, the transfer index *ϕ*(*b, T*\*) equals zero. If the “light side” *p* of *b* equals 2 and *b* is not present in *T*\*, the transfer index *ϕ*(*b, T*\*) equals 1 (because *ϕ*(*b, T*\*) ≤ *p−* 1 [6]). Thus, we do not need to run the *O*(*n*) post-order traversal in this case. We also know that we have already found the minimum transfer distance to *b* during the post-order traversal, if we encounter a branch *b*\* ∈ *T*\* for which *δ*(*b, b*\*) = 1 (because we checked for an exact match before).

### 2.4 Computing Additional Information

The `booster` tool can generate additional TBE-related statistics that might help the user to identify potential problems with his dataset, such as rogue taxa.

This additional statistics include:

–An optional array *A*, storing the taxon transfer index for each taxon. Only BS splits that are “close enough” and reference splits that are “balanced enough” are considered (see below).
–An optional table *B*, storing the percentage of BS trees in which taxon *i* had to be moved between the closest BS split to the reference split *j*. Only BS splits that are “close enough” and reference splits that are “balanced enough” are considered (see below).
–An optional tree *R*, which is a branch-labeled copy of the reference tree, showing branch identifier, depth (the size of the “light side” of the induced split), as well as average transfer distance for each branch.

A split *b*\* ∈ *T*\* is considered closest to a split *b* ∈*T*, if *δ*(*b, b*\*) = *ϕ*(*b, T*\*). Note that there can be multiple closest splits for a split *b*. However, we ensure that we only use one of the possible closest splits for each reference split *b* for calculating the statistics. Let *p* be the size of the “light side” of *b*. Given the user-specified parameter *d* ∈ [0, 1], a closest split *b*\* is considered “close enough” to *b if and only if* 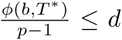 and *b* is considered “balanced enough” *if and only if* 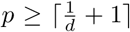. The default value for *d* is 0.3.

The array entry *A*[*i*] is defined as *A*[*i*]:=

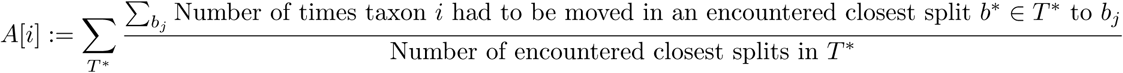

The table entry *B*[*j*][*i*] corresponds to taxon *i* and reference split *b*_*j*_. It is defined as

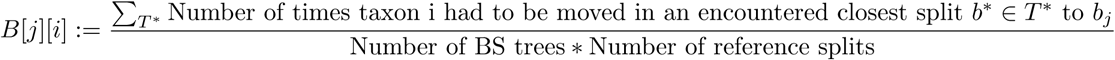

The array *A* and table *B* are based on the computation of “species-to-move” between two bipartitions *b* and *b*\*. The “species-to-move” are a smallest set of taxa that need to be moved to the other side of *b* in order to transform *b* into *b*\*. In case there are multiple such smallest sets, we (as well as `booster`) arbitrarily select one of them for computing the statistics above.

In the `booster` tool, this is computed in linear time by comparing the binary assignment of each taxon.

Our implementation computes “species-to-move” faster by performing an incomplete pre-order traversal, reusing the “number of ones in subtree” values which we already computed during the post-order traversal when searching for the minimum transfer distance. We only descend into a subtree when there is a taxon which needs to be moved in that subtree. Algorithm 3 shows our computation of “species-to-move” via an incomplete pre-order traversal.

## 3 Results

We compared runtime performance and memory consumption between our improved implementation (partially integrated into `RAxML-NG v0.8.1` and later), the naïve implementation previously used in `RAxML-NG v0.7`, and `booster`. Two other popular phylogenetic inference tools also offer TBE computations: `PhyML` [4] and `IQ-Tree` [8]. `IQ-Tree` internally uses `booster` for this task, and `PhyML` can not compute TBE support for user-specified tree sets. Therefore, we excluded `IQ-Tree` and `PhyML` from our evaluation.

### Algorithm 2: Improved *O*(*n* * *m*) Algorithm (used in RAxML-NG v0.8.1 and later)

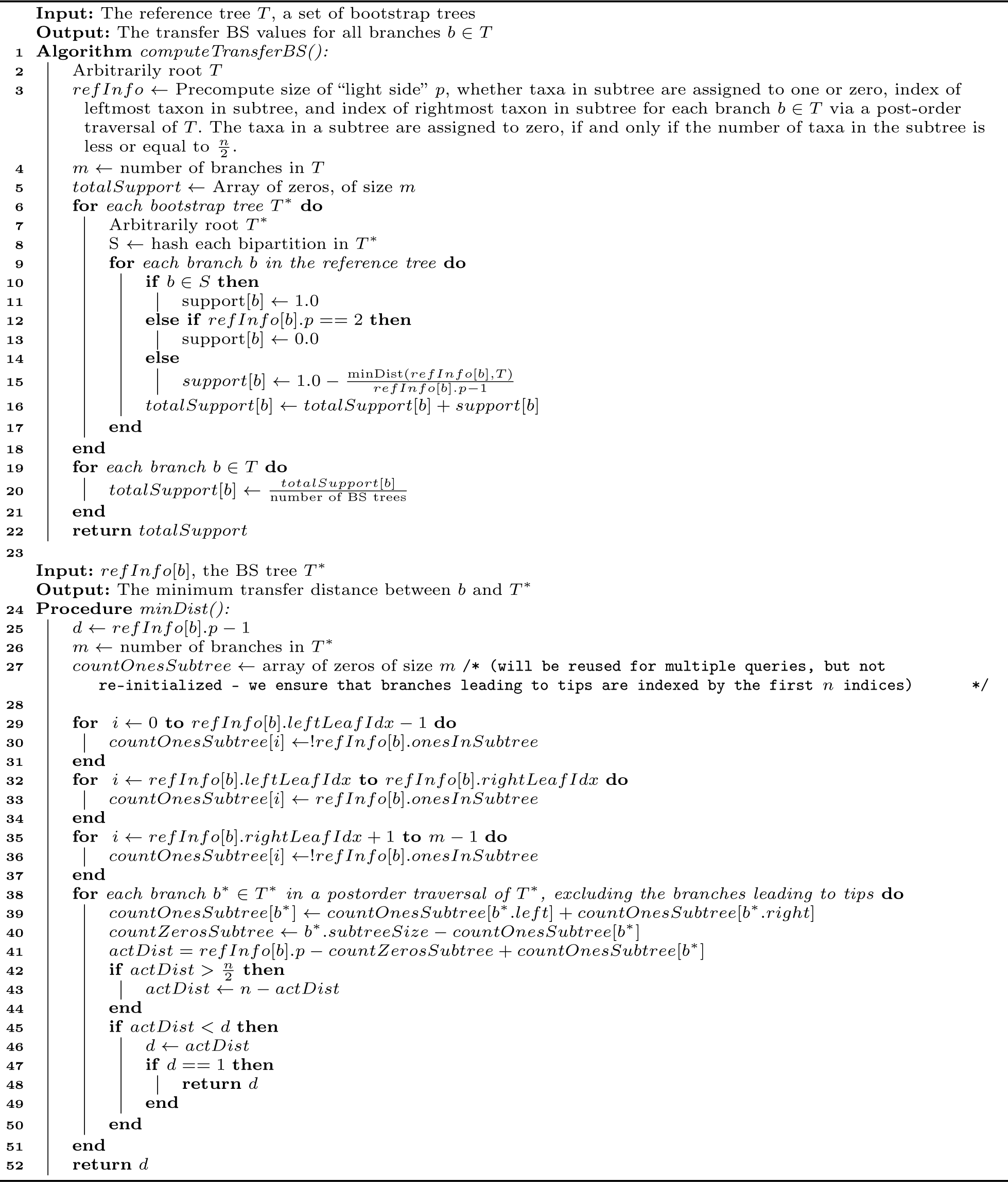

### Algorithm 3: Pre-order traversal to compute “species-to-move” (not integrated yet into the official RAxML-NG repository).

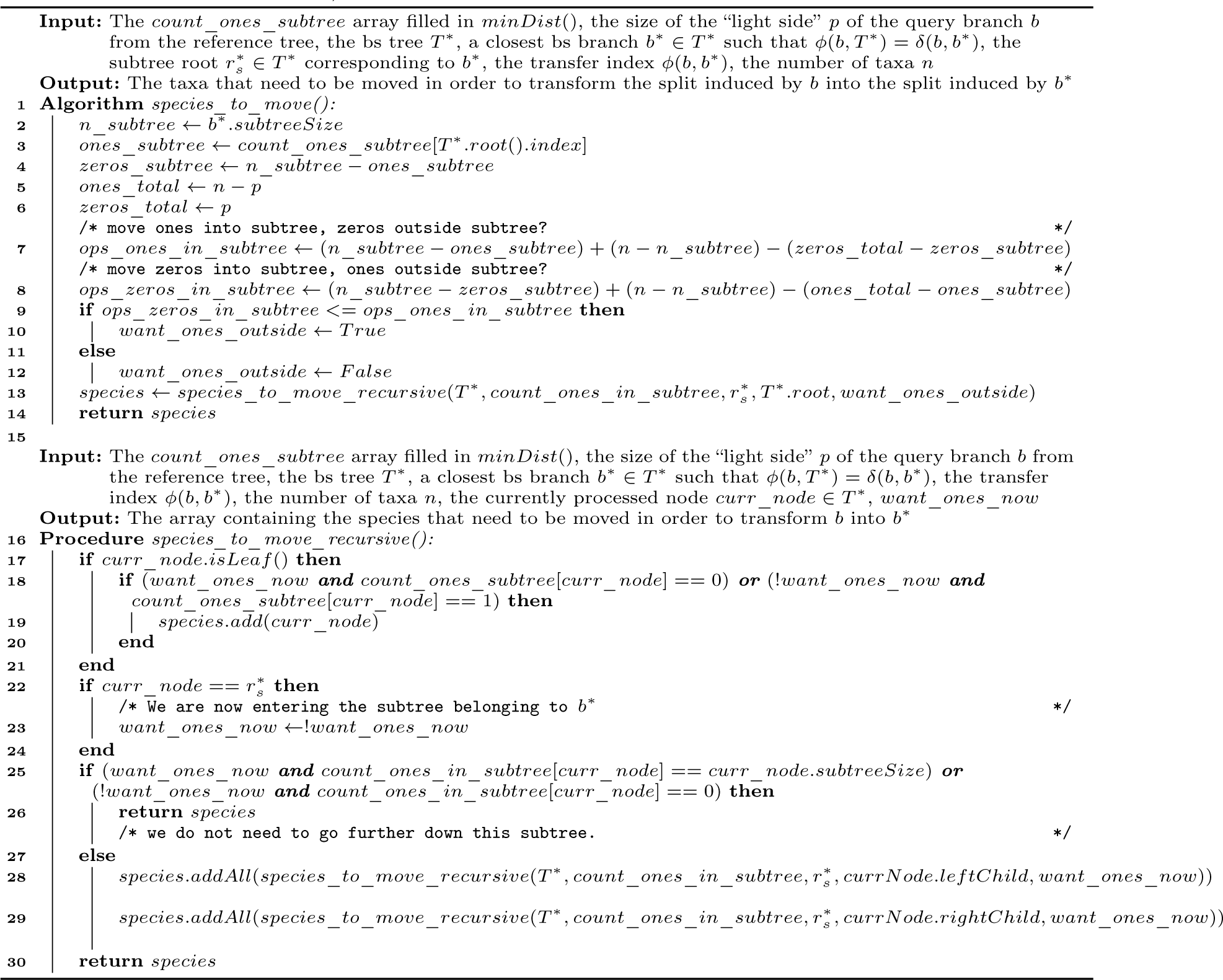

Note that, Truszkowski *et al*. [10] are simultaneously and independently working on an improved algorithm for TBE computations with a lower theoretical run time complexity. The respective prototype implementation is 237 times faster (pers. comm.) than the original `booster` implementation on the dataset C. On this dataset, our implementation is 258 times faster and requires 21 times less memory than `booster`.

We measured runtimes and memory consumption on a machine with two Xeon Gold 6148 (Skylake-SP) CPUs and 768GB RAM. Table 1 lists the empirical datasets we used for evaluation including the average normalized RF distance (nRF) among the BS trees. The nRF shows how similar the BS trees are: e.g., for completely random trees, nRF is close to 1.0. Dataset C is taken from the original TBE study [6]. Datasets A and B were derived from unpublished studies, thus in our supplementary data we anonymized taxon names for these datasets. Datasets D and E were derived from two large 16S rRNA databases, the Living Tree Project v123 (https://www.arb-silva.de/projects/living-tree/) and greengenes (http://greengenes.secondgenome.com/), respectively. Since we lack empirical BS trees for these datasets, we used randomly generated trees instead. The datasets are available at https://doi.org/10.6084/m9.figshare.9692402.

**Table 1.**
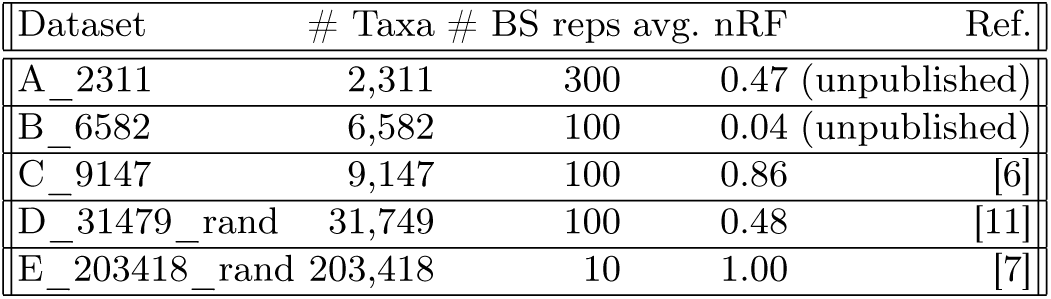
Characteristics of the datasets used for evaluation: number of taxa, number of BS replicates, and average normalized RF distance (nRF) among BS trees.

We used the following tools and tool versions in our performance evaluation:

– `booster` (commit 91fb005 in https://github.com/evolbioinfo/booster/tree/master)
– `RAxML-NG naïve` (commit a4a8f8d in https://github.com/amkozlov/raxml-ng/tree/master)
– `RAxML-NG improved` (commit 757e9be in https://github.com/lutteropp/raxml-ng/tree/tbe)

For each dataset, we measured runtime and memory usage via the command `/usr/bin/time -v`. We averaged the values for `Elapsed (wall clock) time` and `Maximum resident set size` over 3 runs.

Our experimental results show that `RAxML-NG improved` is several orders of magnitude faster than both, `booster`, and `RAxML-NG naïve` on all datasets, while both `RAxML-NG` implementations use considerably less memory than `booster` (see Figures 1 and 2). Despite following different parallelization schemes, both `booster` and `RAxML-NG improved` scale poorly on more than 10 cores (see Figures 3 and 5). However, `booster` shows a drastic increase in required memory whith the number of cores (Figure 4), which is not the case with `RAxML-NG improved` (Figure 6). When computing additional information, both, runtime, and memory usage of `RAxML-NG improved` increase notably, while this is not the case with `booster` (see Figures 7 and 8). In all versions tested, `RAxML-NG improved` showed better runtime performance than `RAxML-NG naïve` and `booster`. Regarding memory usage, both `RAxML-NG` versions required several orders of magnitude less memory than `booster`.

**Fig. 2.**
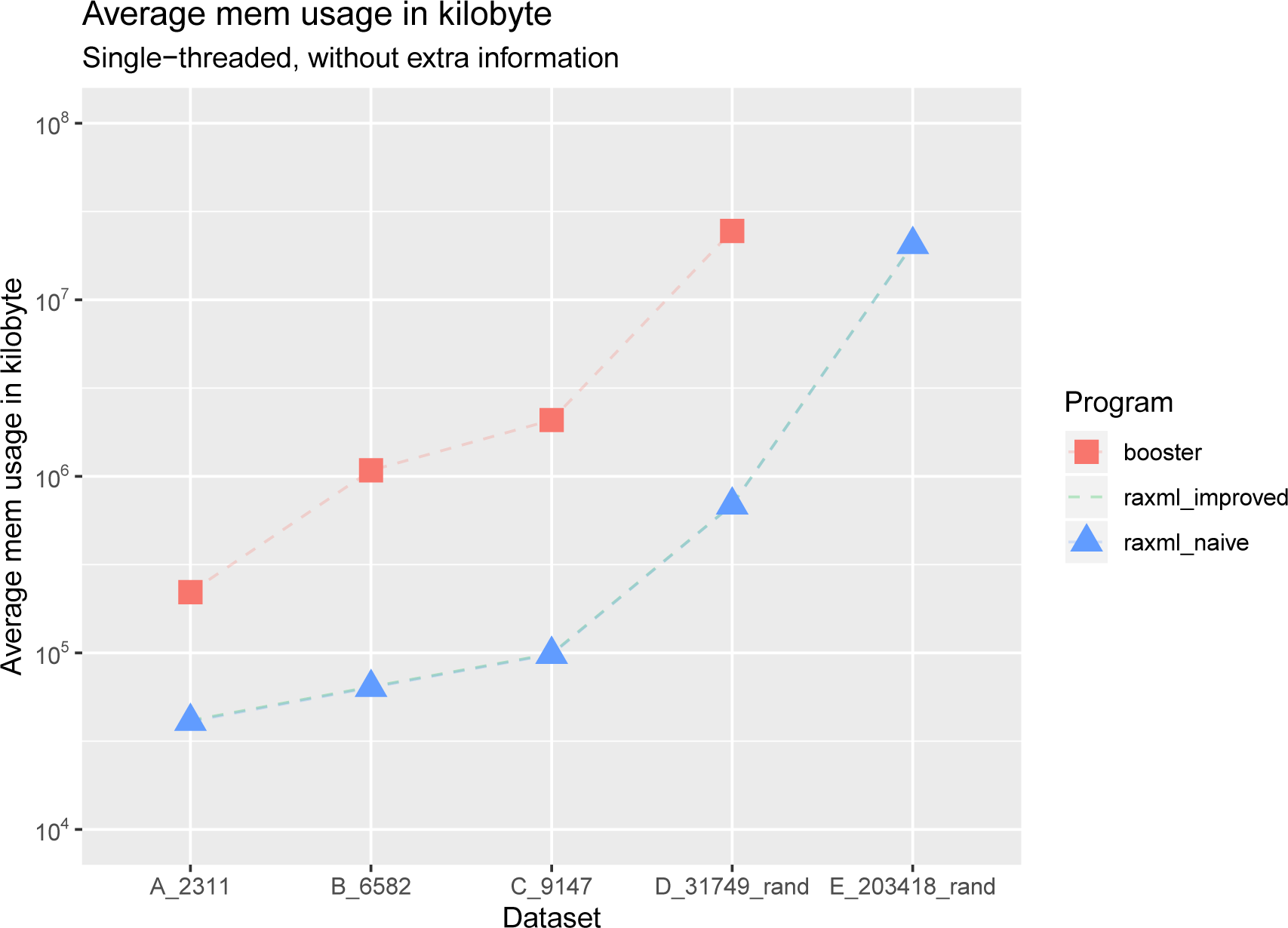
Average total memory usage in kilobytes, without computing additional information. All tools were executed sequentially. Note the logarithmic scale on the y-axis. On the E_203418_rand dataset, `booster` went out of memory. We can see on this plot that `RAxML-NG improved` and `RAxML-NG naïve` require several orders of magnitude less memory than `booster` across all tested datasets. The memory usage of `RAxML-NG improved` and `RAxML-NG naïve` is nearly identical.

**Fig. 3.**
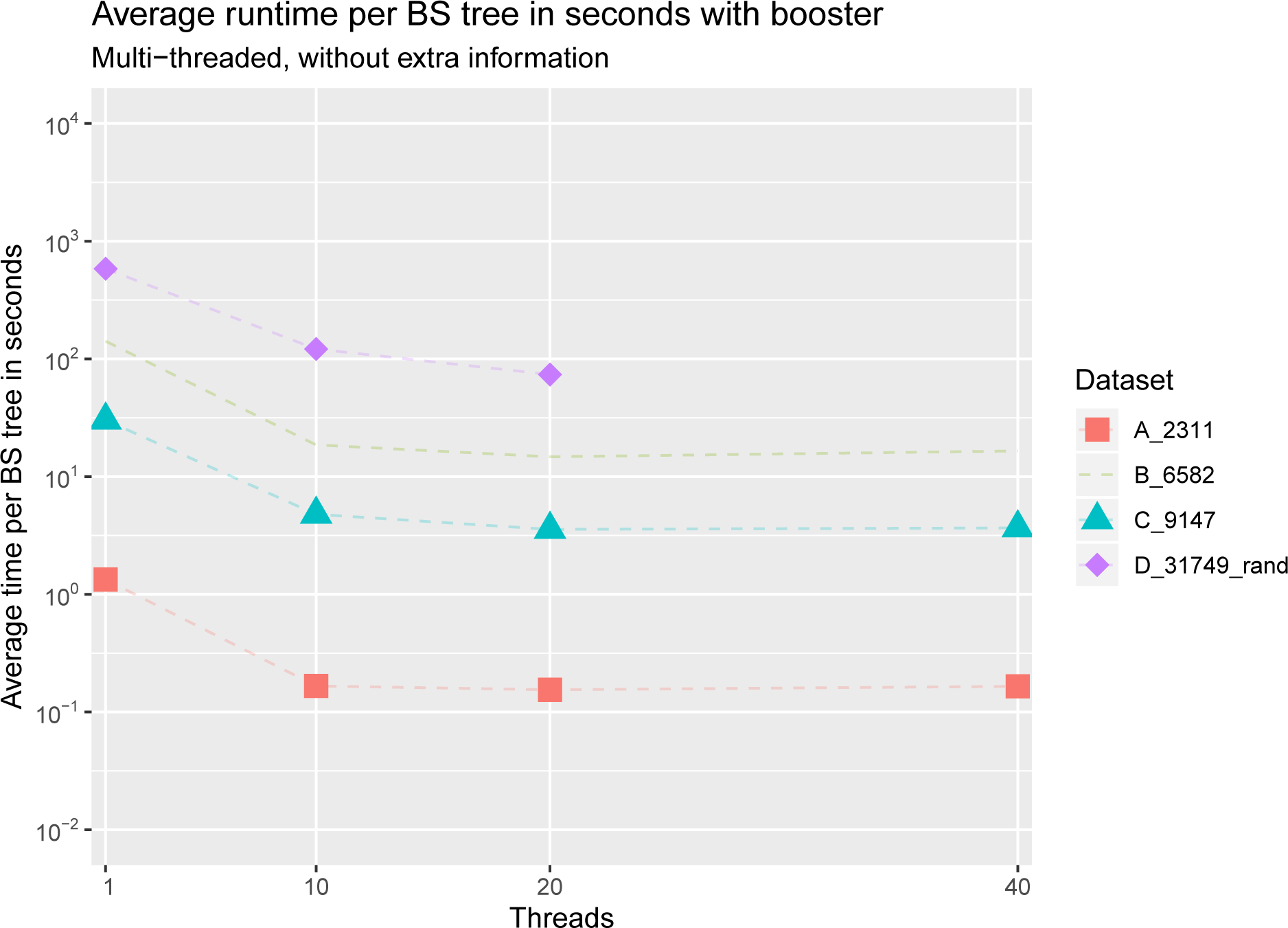
Average runtime per BS tree in seconds for all datasets with `booster`, without computing extra information. Note the logarithmic scale on the y-axis. On the D_31749_rand dataset with 40 threads, `booster` went out of memory. We can see that `booster` does not scale well on more than 10 cores.

**Fig. 4.**
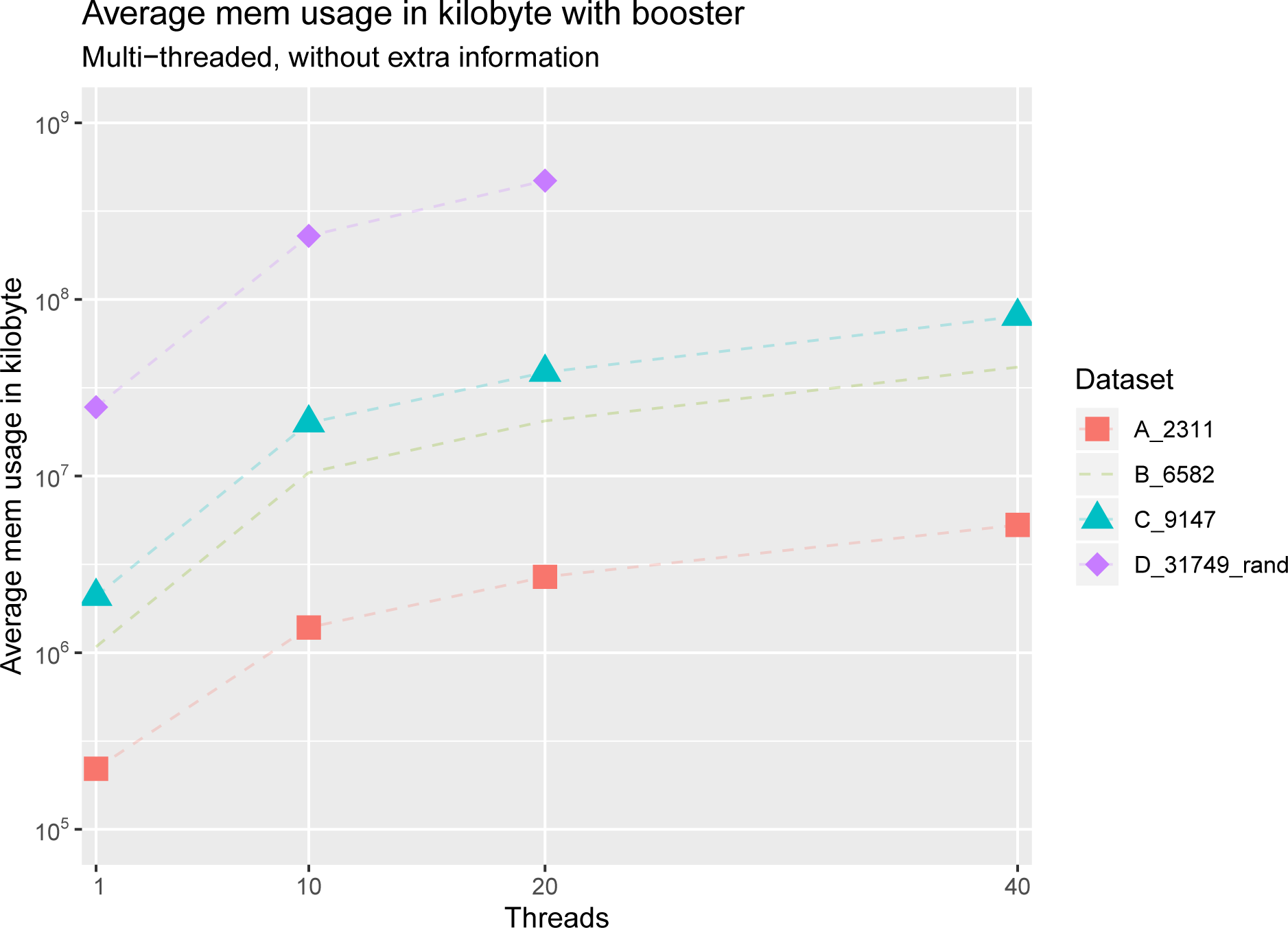
Total memory usage in kilobytes for all datasets with `booster`, without computing extra information. Note the logarithmic scale on the y-axis. On the D_31749_rand dataset with 40 threads, `booster` went out of memory. We can see that the memory consumption of `booster` rises substantially with the number of core.

**Fig. 5.**
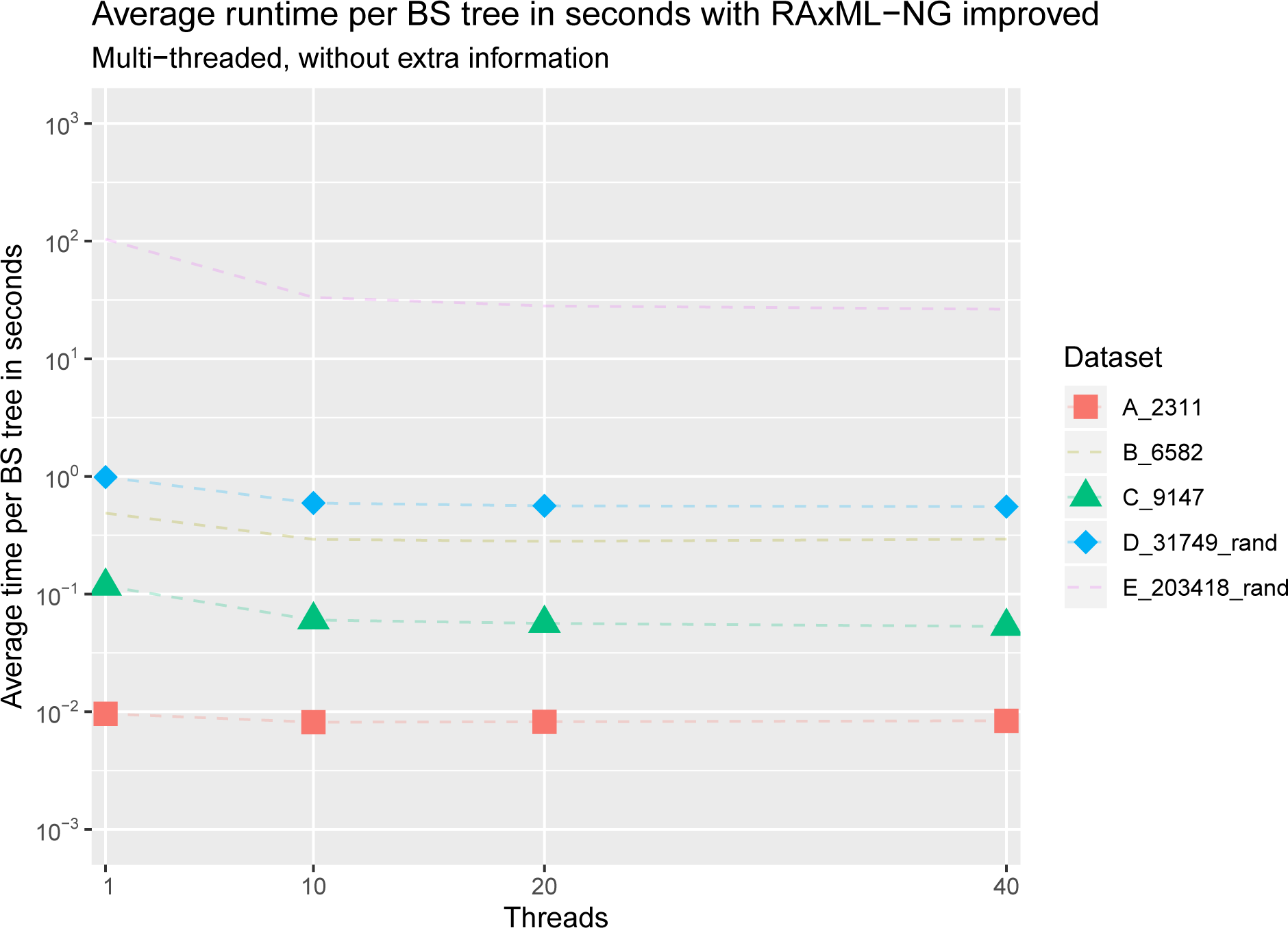
Average runtime per BS tree in seconds for all datasets with `RAxML-NG` improved, without computing extra information. Note the logarithmic scale on the y-axis. We can see that `RAxML-NG improved` does not scale well with more than 10 threads.

**Fig. 6.**
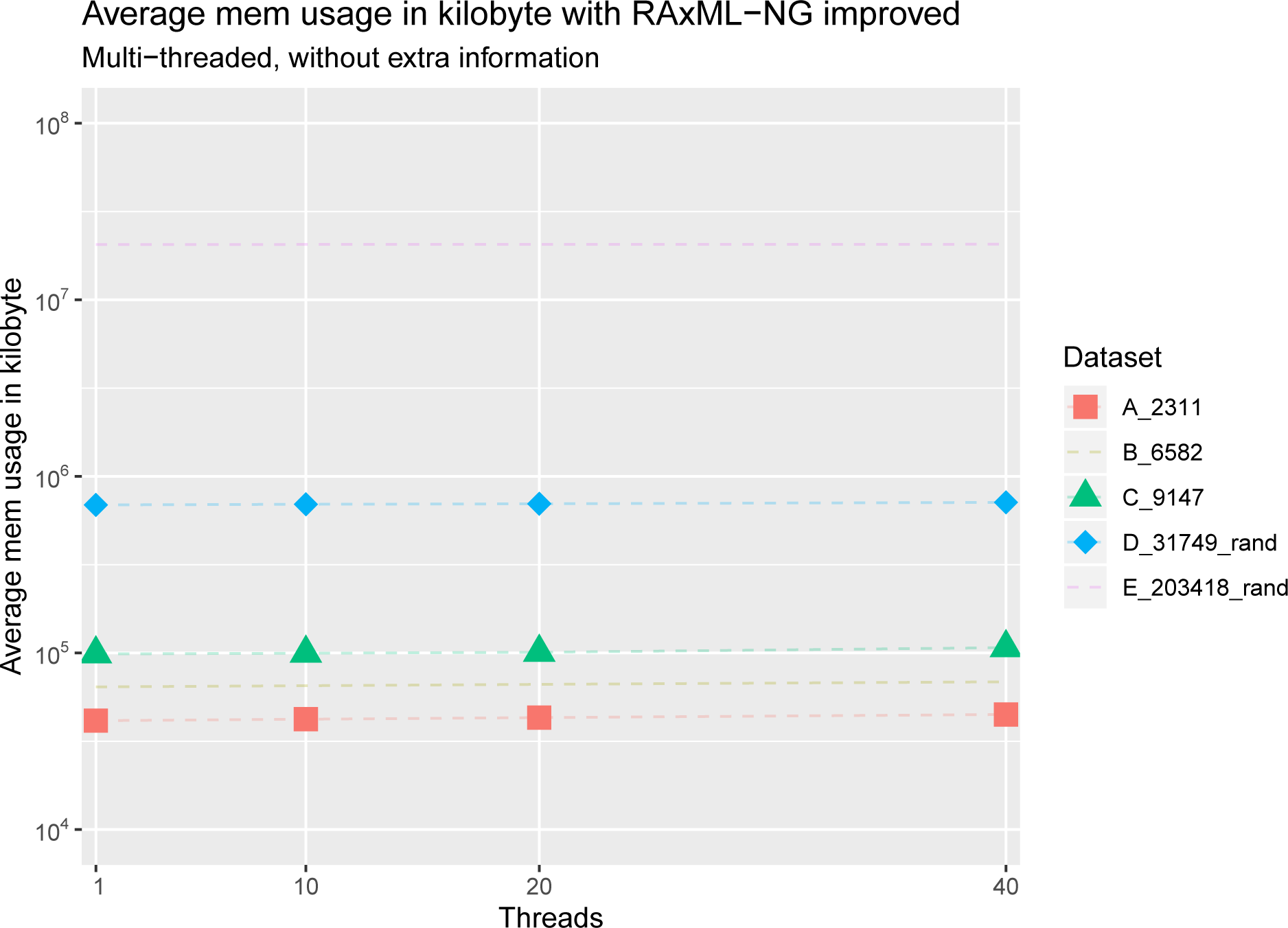
Total memory usage in kilobytes for all datasets with `RAxML-NG improved`, without computing extra information. Note the logarithmic scale on the y-axis. We can see that the memory consumption of `RAxML-NG` improved is not strongly affected by the number of cores.

**Fig. 7.**
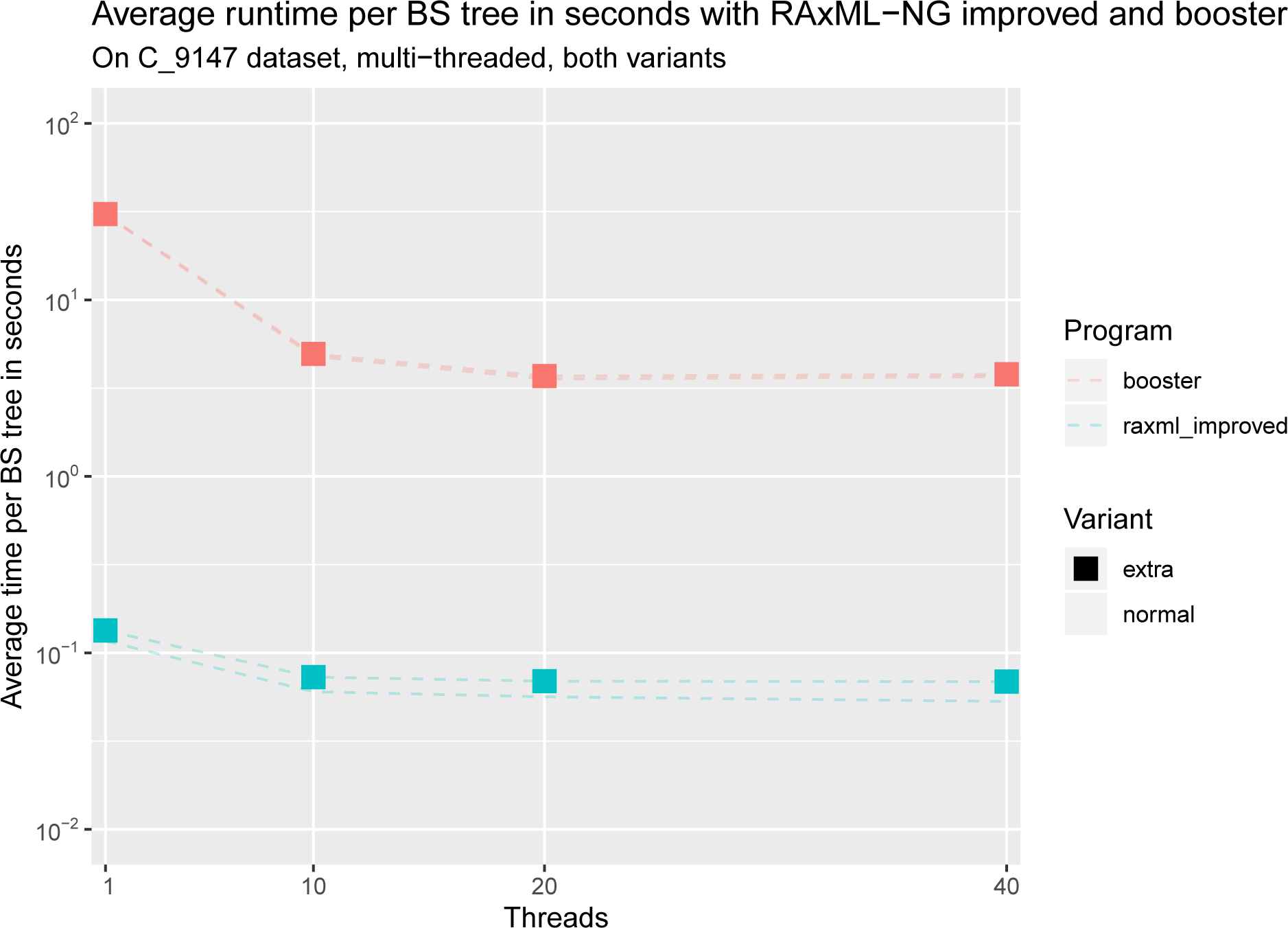
Average runtime per BS tree in seconds for dataset C with `RAxML-NG improved` and `booster`, with and without computing extra information. Note the logarithmic scale on the y-axis. We can see that both tools do not scale well with more than 10 threads and that `RAxML-NG improved` is several orders of magnitude faster than `booster`. Moreover, we can see that the runtimes of `booster` with and without extra information are nearly the same, whereas `RAxML-NG improved` shows an increase in total runtime when computing extra information.

**Fig. 8.**
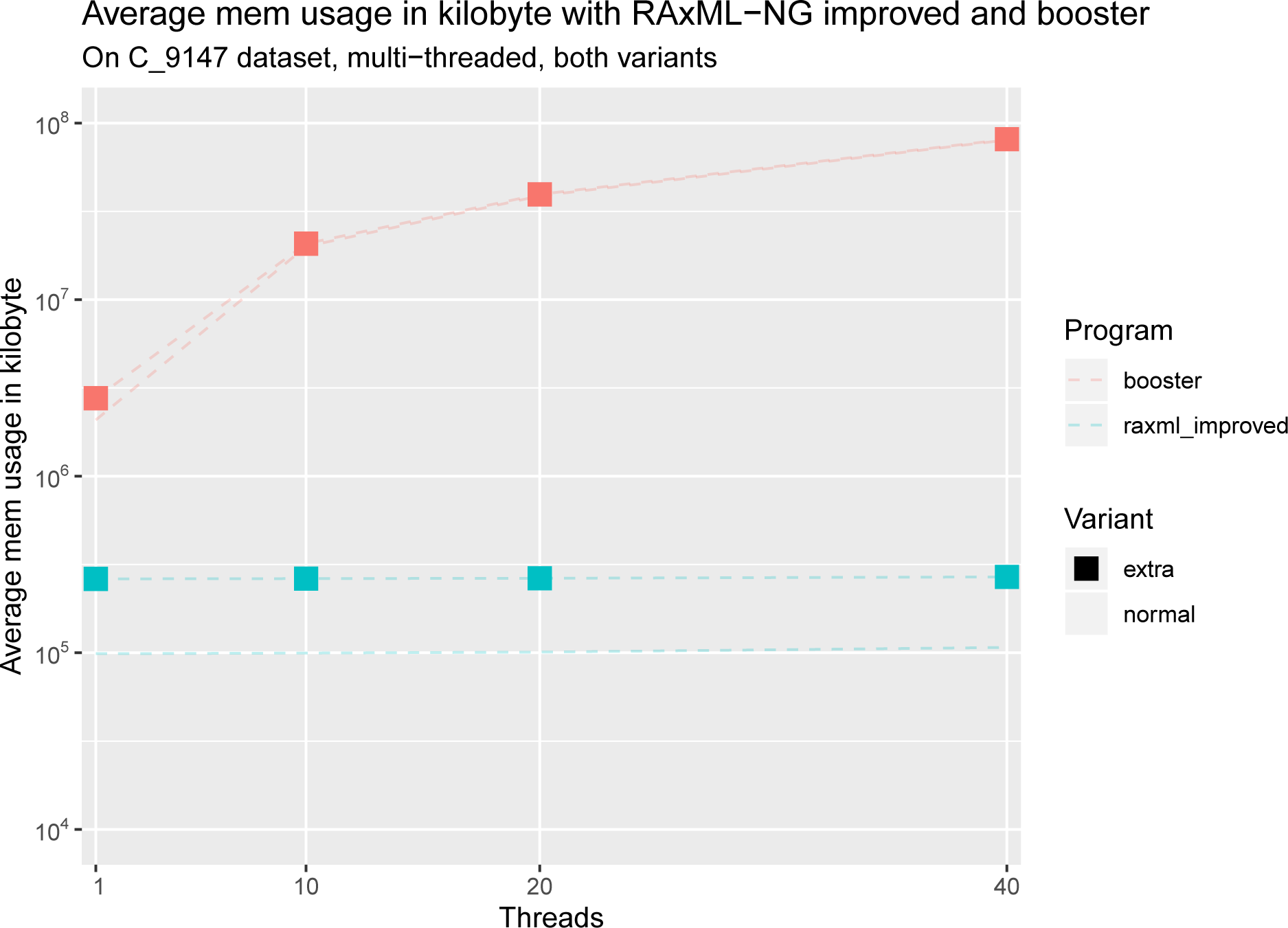
Average total memory usage for dataset C with `RAxML-NG improved` and `booster`, with and without computing extra information. Note the logarithmic scale on the y-axis. We can see that `booster` uses several orders of magnitudes more memory than `RAxML-NG` improved. Moreover, we can see that the memory requirements of `booster` with and without extra information are nearly the same, whereas `RAxML-NG improved` requires more memory when computing extra information.

## 4 Conclusions

We developed and made available a substantially faster and more memory-efficient Transfer Bootstrap implementation, which allows to calculate TBE support metrics on extremely taxon-rich phylogenies, without constituting a computational limitation. For example, on dataset D with 31, 749 taxa and 100 BS replicates using a single thread, our implementation `RAxML-NG improved` computed TBE support values in under two minutes, while `RAxML-NG naïve` and `booster` required 458 minutes and 916 minutes, respectively.

## 5 Future Work

Instead of selecting only one possibility for determining “species-to-move” to transform one bipartition into another, one could average over all possible minimal sets of taxa when computing the additional statistics. As most entries in the extra table *B* are zero, memory usage could be further reduced by using a sparse matrix representation and changing the output format to only print nonzero cells. Our parallelization could be improved by also parallelizing across BS trees or using MPI to orchestrate transfer bootstrap computations onto multiple cluster nodes.

## Acknowledgement

We thank Frédéric Lemoine (the author of the `booster` tool) for the helpful discussions and for his valuable feedback on this manuscript. Part of this work was funded by the Klaus Tschira foundation.

## References

1. Brehelin, L., Gascuel, O., Martin, O.: Using repeated measurements to validate hierarchical gene clusters. Bioinformatics 24(5), 682–688 (2008)

2. Felsenstein, J.: Confidence limits on phylogenies: an approach using the bootstrap. Evolution 39(4), 783–791 (1985)

3. Flouri, T., Izquierdo-Carrasco, F., Darriba, D., Aberer, A.J., Nguyen, L.T., Minh, B., Von Haeseler, A., Stamatakis, A.: The phylogenetic likelihood library. Systematic biology 64(2), 356–362 (2014)

4. Guindon, S., Dufayard, J.F., Lefort, V., Anisimova, M., Hordijk, W., Gascuel, O.: New algorithms and methods to estimate maximum-likelihood phylogenies: assessing the performance of phyml 3.0. Systematic biology 59(3), 307–321 (2010)

5. Kozlov, A.M., Darriba, D., Flouri, T., Morel, B., Stamatakis, A.: RAxML-NG: a fast, scalable and user-friendly tool for maximum likelihood phylogenetic inference. Bioinformatics (05 2019). https://doi.org/10.1093/bioinformatics/btz305, https://doi.org/10.1093/bioinformatics/btz305

6. Lemoine, F., Entfellner, J.B.D., Wilkinson, E., Correia, D., Felipe, M.D., Oliveira, T.d., Gascuel, O.: Renewing felsenstein’s phylogenetic bootstrap in the era of big data. Nature 556(7702), 452 (2018)

7. McDonald, D., Price, M.N., Goodrich, J., Nawrocki, E.P., DeSantis, T.Z., Probst, A., Andersen, G.L., Knight, R., Hugenholtz, P.: An improved Greengenes taxonomy with explicit ranks for ecological and evolutionary analyses of bacteria and archaea. The ISME journal 6, 610-8 (2012). https://doi.org/10.1038/ismej.2011.139

8. Nguyen, L.T., Schmidt, H.A., von Haeseler, A., Minh, B.Q.: Iq-tree: a fast and effective stochastic algorithm for estimating maximum-likelihood phylogenies. Molecular biology and evolution 32(1), 268–274 (2014)

9. Pattengale, N.D., Alipour, M., Bininda-Emonds, O.R., Moret, B.M., Stamatakis, A.: How many bootstrap replicates are necessary? Journal of Computational Biology 17(3), 337–354 (2010)

10. Truszkowski, J.M., Gascuel, O., Swenson, K.: Rapidly computing the phylogenetic transfer index. bioRxiv (2019). https://doi.org/10.1101/743948, https://www.biorxiv.org/content/early/2019/08/22/743948

11. Zanne, A.E., Tank, D.C., Cornwell, W.K., Eastman, J.M., Smith, S.A., FitzJohn, R.G., McGlinn, D.J., O’Meara, B.C., Moles, A.T., Reich, P.B., et al.: Three keys to the radiation of angiosperms into freezing environments. Nature 506(7486), 89 (2014)

